# An RXLR effector targets ER-Golgi interface to induce ER stress and necrotic cell death

**DOI:** 10.1101/2023.12.15.571945

**Authors:** Jihyun Kim, Jesse Kaleku, Jongchan Woo, Hongshi Jin, Hui Jeong Kang, Minji Kang, Haeun Kim, Seungmee Jung, Cecile Segonzac, Eunsook Park, Doil Choi

**Author notes:** The authors responsible for distribution of materials integral to the findings presented in this article in accordance with the policy described in the Instructions for Authors (https://academic.oup.com/plcell/pages/General-Instructions) are: Eunsook Park, Doil Choi.

## Abstract

To achieve successful colonization, the pathogen secretes hundreds of effectors into host cells to manipulate the host’s immune response. Despite numerous studies, the molecular mechanisms underlying effector-induced necrotic cell death remain elusive. In this study, we identified a novel virulent RXLR effector named Pc12 from *P. capsici.* Pc12 induces necrosis by triggering a distinct ER stress response through its interaction with Rab13-2. Unlike conventional hypersensitive response cell death associated with effector-triggered immunity, Pc12-induced cell death does not coincide with defense gene expression. Instead, it induces the aggregation of ER-resident proteins and confines secretory proteins within the ER. Pc12 interacts with Rab13-2, exhibiting a specific affinity for the active form of Rab13-2. Thus, the complex of Pc12 and Rab13-2 mimics the conformation of the inactive state of Rab13-2, subsequently recruiting the Rab-escort protein (REP). This process results in disruptions in vesicle formation within the ER-Golgi trafficking pathway. Furthermore, the substitution of a single amino acid of Rab13-2 structurally predicted to be crucial for the Pc12 interaction decreased the interaction with Pc12 while maintaining the interaction with REP1. These findings offer valuable insights into the ER stress-mediated cell death as well as a potential strategy for enhancing resistance against pathogens.

## Introduction

*P. capsici* is an oomycete pathogen causing blight disease in economically important crops, including Solanaceae, Cucurbitaceae, Fabaceae, and Malvaceae (Kamoun et al. 2015; Parada-Rojas et al. 2021; Quesada-Ocampo et al. 2023). *P. capsici* can inflict up to 100% crop damage in fields upon successful plant infection. This is attributed to its rapid proliferation under various environmental conditions, robust persistence in the soil over five years, and high genetic variability, leading to its classification as the fifth most destructive oomycete worldwide (Kamoun et al. 2015; Quesada-Ocampo et al. 2023). *Phytophthora* species secrete an arsenal of RXLR effector proteins to infect host plants. RXLR effectors feature a highly conserved Arg-X-Leu-Arg (RXLR) motif, which is essential for their transport into host cells (Whisson et al. 2007; Wawra et al. 2017). They play a significant role in facilitating the successful establishment of the pathogen by disrupting plant physiology and defense responses (Whisson et al. 2007; Fan et al. 2018; Boevink et al. 2020). Therefore, a comprehensive understanding of the role of RXLR effectors is important for devising strategies to manage plant diseases effectively.

The endoplasmic reticulum (ER) functions as a cellular factory responsible for protein synthesis, folding, and export to subcellular organelles. These sequential processes must be balanced to maintain cellular homeostasis under environmental conditions (Cao and Kaufman 2012; Angelos et al. 2017; Adams et al. 2019). However, when an imbalance occurs leading to the accumulation of misfolded or unfolded proteins, known as ER stress, eukaryotic cells initiate the unfolded protein response (UPR). The UPR involves the translocation of key transcription factors like bZIP28 and bZIP60 into the nucleus via maturation processes, such as proteolytic cleavage or appropriate splicing. These transcription factors regulate the expression of UPR target genes crucial for protein folding and ER-associated degradation (ERAD), enhancing protein folding capacity in cells (Cao and Kaufman 2012; Angelos et al. 2017; Adams et al. 2019). However, if plants are subjected to severe or chronic stress, it can further lead to unresolved ER stress-induced programmed cell death (PCD) as a survival mechanism (Yang et al. 2014; Liu and Howell, 2016; Yang et al. 2021; Simoni et al. 2022).

Rab is a superfamily of small GTPases that plays a crucial role in intracellular vesicle trafficking, an essential process for the transfer of proteins, lipids, and other cellular materials between different membrane compartments within eukaryotic cells (Nielsen et al. 2008; Stenmark et al. 2009; Pylypenko et al. 2018; Gray et al. 2020). Within a species, Rab proteins exist in multiple isoforms, each defining membrane identity and mediating vesicle trafficking to distinct subcellular compartments. Rab proteins function in the formation and guide of vesicles by acting as molecular switches, cycling between an active GTP-bound form and an inactive GDP-bound form (Nielsen et al. 2008; Stenmark et al. 2009; Pylypenko et al. 2018; Gray et al. 2020). Dysfunctional Rab proteins in vesicle trafficking can result in ER stress due to improper protein compartmentation (Kiral et al., 2018). Several effectors of various pathogens are known to suppress antimicrobial protein secretion (PR1 and PDF1.2) or subvert vesicle movement toward the pathogen’s focal area by targeting Rab proteins (Tomczynska et al. 2018; Pandey et al. 2021; Li et al. 2022). However, it remains elusive how the RXLR effector leads to ER stress-induced cell death by disrupting Rab proteins in vesicle trafficking.

In this study, we report a virulent RXLR effector Pc12 from *P. capsici*, triggering necrotic cell death in plants of the Solanaceae. Transient overexpression and host-induced gene silencing (HIGS) of *Pc12* in plants significantly affected the pathogenicity of *P. capsici*. Pc12 induced ER stress, resulting in distinct expression patterns of UPR genes and altering the location of ER-resident proteins and the secretory pathway. Notably, in the presence of Pc12, an active Rab13-2 attracted the Rab-escort protein 1 (REP1), enhancing its interaction. We have also identified a crucial residue in Rab13-2 to avoid being targeted by Pc12 while maintaining its original function to interact with REP1 and PRA1. Consequently, this study provides valuable insights suggesting that modifying Rab13-2 to evade effector targeting could offer a promising strategy for combating the pathogen infection while preserving its essential functions.

## Results

### Pc12 induces cell death independently of the plant defense mechanism in Solanaceae plants

Among the previously selected RXLR effector candidates (Seo et al. 2023), Pc12 exhibited a strong induction of cell death in *N. benthamiana*. Interestingly, Pc12 consistently induces cell death in other Solanaceae species when transiently expressed in *N. tabacum, C. annuum,* and *S. lycopersicum* (Figure 1A), suggesting that Pc12-induced cell death is more pronounced within the Solanaceae family. To determine whether the observed cell death induced by Pc12 resulted from intracellular receptors of nucleotide-binding site leucine-rich repeats (NLRs) - mediated hypersensitive response (HR), we investigated Pc12-mediated cell death in *N. benthamiana* plants with individual silencing of NLR chaperon (*SGT1*) or NLR-downstream signaling factors, including *EDS1, ADR1/NRG1,* and *NRC234* (Botër et al. 2007; Dongus and Parker, 2021; Wu et al. 2017). Interestingly, Pc12-induced cell death remained unaffected by the silencing of NLR chaperon or NLR-downstream signaling components (Figure 1B-D), suggesting that Pc12 induces cell death through mechanisms that are distinct from those typically associated with NLR-mediated effector-triggered immunity (ETI). To further validate whether the cell death induced by Pc12 is distinct from HR cell death, we compared the expression of defense-related genes during Pc12 expression in contrast to the expression of a bacterial effector derived from *Xanthomonas spp.,* XopQ. XopQ is known to interact with the resistance protein Roq1 in *N. benthamiana*, resulting in a mild chlorotic phenotype (Schultink et al. 2017). Transcription of defense-related genes, including *PR1, RbohB,* and *WRKY8*, exhibited significant upregulation upon transient overexpression of XopQ. In contrast, Pc12 expression did not significantly induce the expression of defense-related genes (Figure 1E), supporting the idea that Pc12-induced cell death may activate a different molecular pathway compared to the ETI-induced HR cell death.

**Figure 1.**
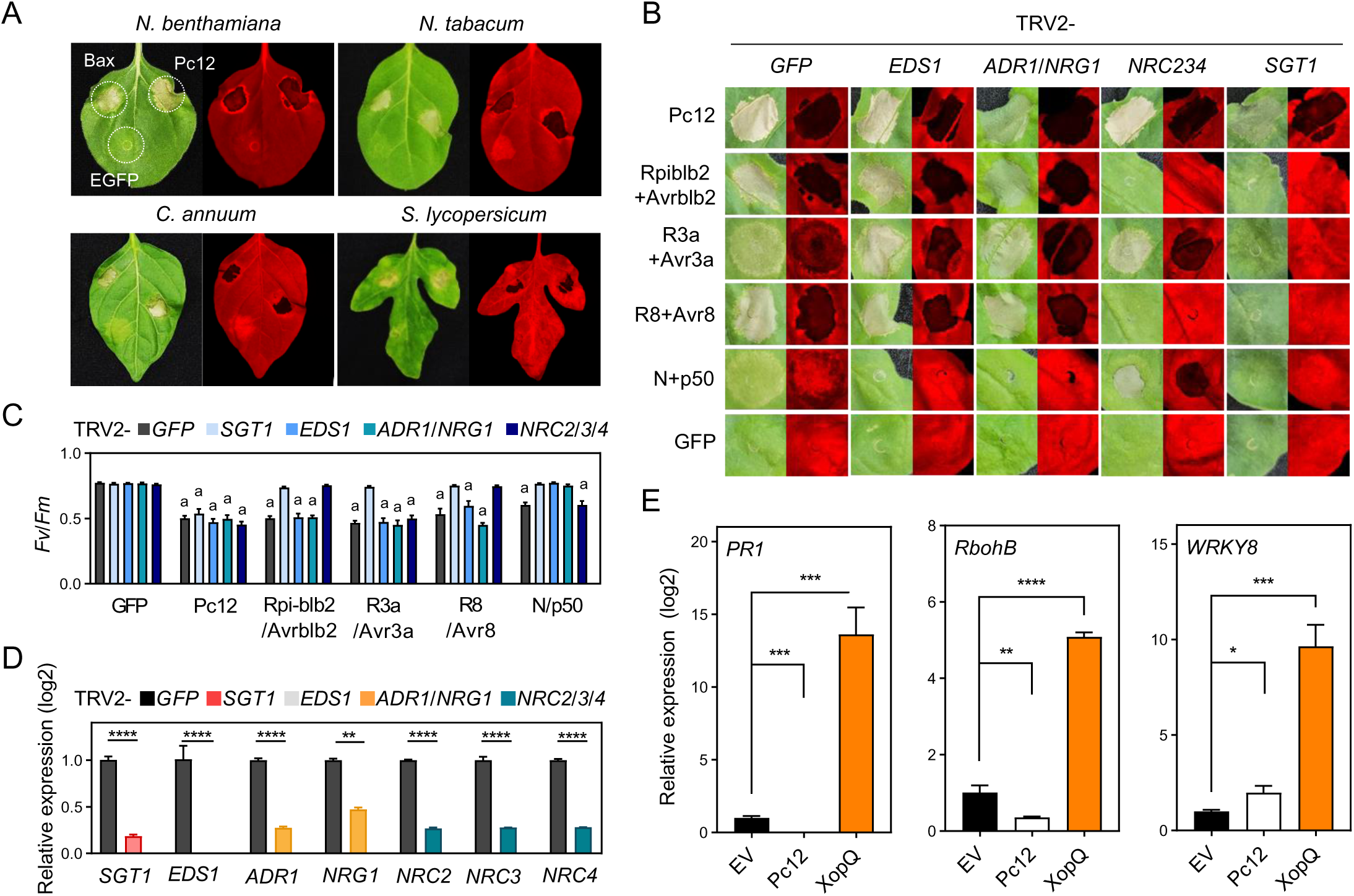
Pc12 causes cell death independent of defense response in Solanaceae. (A) Pc12-triggered cell death in *Nicotiana benthamiana, Nicotiana tobacum, Capsicum annuum*, and *Solanum lycopersicum*. Bax and EGFP were used as a positive control and a negative control, respectively. The leaves of 4-week-old of plants were agroinfiltrated. The images were photographed under white and UV light at 2 days after agroinfiltration. (B) Cell death induced by Pc12 in *N. benthamiana* with silenced NLR-downstream signaling genes *EDS1*, *ADR1/NRG1*, *NRC2/3/4*, and *SGT.* Positive controls consisted of the combinations Rpiblb2+Avrblb2, R3a+Avr3a, R8+Avr8, and N+p50, while GFP served as the negative control. Agrobacterium carrying each construct was infiltrated into the leaves of 5-week-old plants with the respective gene components silenced, and images were taken at 2 dpi. (C) Cell death observed in images (B) was quantified using the quantum yield (*Fv/Fm*) using a closed FluorCam system. The data represents the mean ± standard deviation (SD, n = 7-10). a indicates statistically significant differences by Student t-test (****, P < 0.0001). (D) qRT-PCR analysis of the transcripts level in *N. benthamiana*-silencing *SGT1, EDS1, ADR1/NRG1*, and *NRC2/3/4.* Leaf disks were sampled at 5-week-old in each components-silenced *N. benthamiana.* Asterisks denote statistically significant differences by Student t-test (****, P < 0.0001). Data are mean ± SD. (E) Expression patterns of *PR1, RbohB*, and *WKRY8* in *N. benthamiana* expressing Pc12, XopQ, and empty vector (EV). Total RNA was extracted in 30 hour-post-infiltration. Asterisks denote statistically significant differences by Student t-test (*, P < 0.1; **, P < 0.01; ***, P < 0.001; ****, P < 0.0001). Data are mean ± SD. (3 experimental repeats).

### Pc12 functions as a virulence effector, enhancing the pathogenicity of *P. capsici*

*Pc12* expression is elevated during the biotrophic phase of *P. capsici* (Supplemental Figure 1). Although Pc12 displays the canonical structure of an RXLR effector, it remains uncertain whether Pc12 is a virulent effector for *P. capsici* colonization. To assess the impact of Pc12 on the virulence of *P. capsici*, the pathogen was inoculated in plants expressing Pc12 under the control of an ethanol-inducible promoter without ethanol treatment (Figure 2A). Leaky expression of Pc12 driven by the ethanol-inducible promoter (Lee et al. 2018) allows us to generate weak expression of Pc12 avoiding rapid cell death. The *Agrobacterium* carrying construct of RFP or RFP-Pc12 was infiltrated into *N. benthamiana*, followed by drop inoculation of *P. capsici* onto the leaves 1 day post infiltration. We observed a substantial increase in lesion size (Figure 2A) and *P. capsici* biomass (Figure 2B). Host-induced gene silencing (HIGS) is a method that initiates targeted gene suppression in the pathogen through the host’s RNA interference (RNAi) machinery (Cheng et al. 2022). In all cases, the silencing of Pc12 by HIGS using TRV resulted in reduction in the pathogenicity of *P. capsici* compared to the control (Figure 2C-D). This suggests that Pc12 plays a role in enhancing the pathogenicity of *P. capsici* acting as a virulence effector.

**Figure 2.**
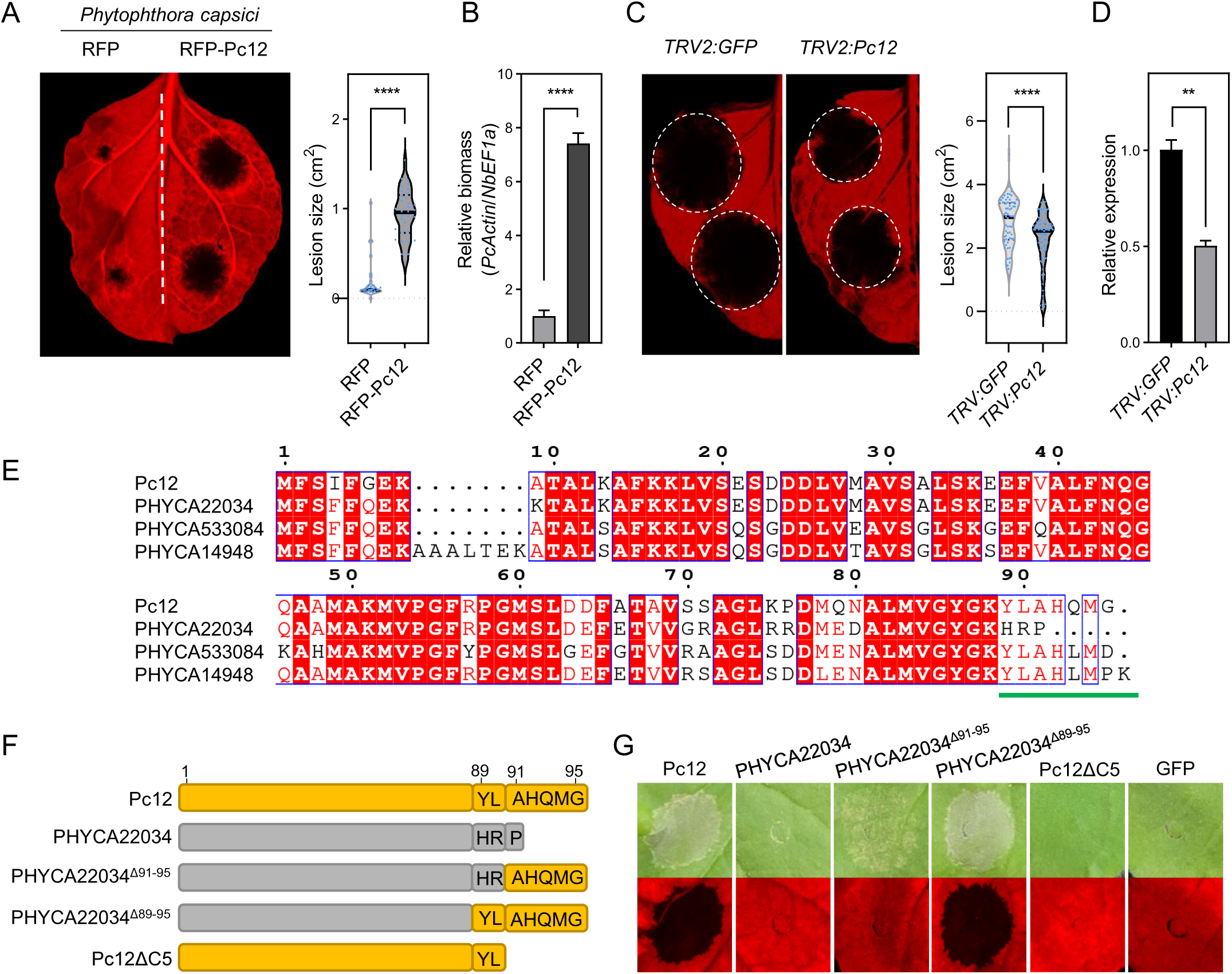
Pc12 enhances the growth of *P. capsici*, triggering necrotic cell death through its C-terminus. (A) Increased growth of *P. capsici* under Pc12 expression compared to a control. RFP and RFP-Pc12 were expressed by ethanol-inducible promoter in *N. benthamiana*. *P. capsici* was inoculated on the leaves at 1 day-post-agroinfiltration without ethanol treatment. Images were taken under UV light at 2 day-post-inoculation. The lesion sizes were measured using ImageJ. Asterisks denote statistically significant differences using a Student t-test (****, P < 0.0001). Data are mean ± SD. (B) Relative biomass of *P. capsici* in the leaves shown in Figure 2A. Leaf disks from around the inoculated area were sampled at 2 dpi. Total genomic DNA was extracted and subjected to qPCR analysis. The *P. capsici* biomass was quantified by the *PcActin* normalized to the *NbEF1α*. Asterisks denote statistically significant differences by Student t-test (****, P < 0.0001). Data are mean ± SD. (C) Suppression of *P. capsici* symptoms through Pc12 silencing via HIGS. Plants were agroinfiltrated with TRV carrying GFP and Pc12. At 14 dpi, *P. capsici* was inoculated on the upper leaves. Images were taken at 2 dpi, and the lesion size was measured by ImageJ. (D) Relative expression of Pc12 in Pc12-silenced and GFP leaves. The inoculated leaves were sampled at 6 hpi and utilized in qRT-PCR analysis. The transcripts level of Pc12 was normalized to the *PcTubulin*. Asterisks denote statistically significant differences by Student t-test (**, P < 0.01; ****, P < 0.0001). Data are mean ± SD. (E) Pairwise sequence alignment comparisons of Pc12 homologs of *P. capsici* genome (LT1534). Alignments were obtained using the MUSCLE algorithm and were visualized via ESPript 3.0 (Robert and Gouet 2014). Strictly or highly conserved residues are highlighted in red boxes or blue empty boxes, respectively. (F) Schematic representation of C-terminal chimeras between Pc12 and Pc22034. (G) Cell death induced by the Pc12, Pc22034, Pc22034 mutants, and Pc12ΔC5 described in (F)

### Pc12 lineage expanded specifically in *P. capsici* and the C-terminal residue is crucial for inducing cell death

To elucidate whether Pc12 functions as a single gene or as part of a gene family, we examined the copy number variation and sequence polymorphism of Pc12 homologs in *Phytophthora spp*. The sequence polymorphism of Pc12 was assessed through a BLAST search, encompassing genome sequences from *P. capsici* (ten strains; LT1534, KPC-7, MY-1, JHAI1-7, and CPV-219/262/267/270/277/302) and four other *Phytophthora spp.* (*P. ramorum* Pr102, *P. sojae* P6497, *P. cinnamoni* CBS144.22, and *P. infestans* T30-4) retrieved from the public database. A comparative genomic analysis revealed that the *P. capsici* strains exhibited a significantly higher number of Pc12 homologs in their genome compared to other *Phytophthora* pathogens (Supplemental Table 1, Reyes-Tena et al. 2019; NCBI reference sequence database (RefSeq)). In order to ascertain whether the Pc12 homologs of these *Phytophthora spp.* could induce cell death, three homologs with high similarity to Pc12, except for 90% identity with Pc12, from *P. capsici* LT1534 and one homolog each from *P. ramorum, P. sojae,* and *P. cinnamomi* were synthesized and transiently expressed in *N. benthamiana*. Interestingly, the cell death phenotype was only observed with *P. capsici* homologs, with the exception of PHYCA22034 (Supplemental Figure 2). Although Pc12 and PHYCA22034 share 85% identity, substantial differences are found at the C-terminal end, which may be responsible for their distinct abilities to induce cell death (green line in Figure 2E). To determine the importance of the C-terminus of Pc12 in triggering cell death, we generated a series of chimeric mutants with modified amino acids at the C-terminus of PHYCA22034 and a deletion mutant lacking five amino acids of Pc12 (Pc12ΔC5) (Figure 2F). The substitution of amino acids from the 91st to 95th position of Pc12 in PHYCA22034 resulted in mild cell death, whereas the substitution of amino acids from the 89th to the 95th position fully restored cell death in *N. benthamiana* (Figure 2G). Consistently, Pc12ΔC5 reduced the induction of cell death (Figure 2G). These results suggest that the functional Pc12 lineage has expanded specifically within *P. capsici*. Furthermore, the C-terminal sequence plays a pivotal role in triggering cell death in plants.

### Pc12 interaction with Rab13-2 small GTPase via C-terminal five amino acids is essential for cell death

To identify Pc12 interacting proteins that would be responsible to induce necrotic cell death, we performed immunoprecipitation and mass spectrometry (IP-MS) using Pc12 and Pc12ΔC5. The list of proteins exclusively interacting with Pc12 but not with Pc12ΔC5 includes a small GTPase Rab13-2 protein in the highest peptide-spectrum match (PSM) score (Supplemental Table 2). We confirmed *in planta* interaction between Pc12 and Rab13-2 by a co-immunoprecipitation (co-IP) assay (Figure 3A). In addition, homologs of Pc12 that trigger cell death also presented interaction with Rab13-2 (Supplemental Figure 2 and Figure 3A). This suggests that the interaction between Pc12 homologs, which share a similar C-terminus with Rab13-2, is crucial for the induction of cell death. Rab small GTPase is a superfamily protein functionally conserved in eukaryotic cells (Nielsen et al. 2008, Pylypenko et al. 2018, Nielsen et al. 2020). Typically, the family comprises more than 60 isoforms that reside in specific membrane compartments generated during membrane trafficking in eukaryotic cells (Stenmark 2009, Pylypenko et al. 2018, Nielsen et al. 2020). In the *N. benthamiana* genome, the Rab protein family consists of 149 genes (Supplemental Figure 3). To explore the specificity of the interaction between Pc12 and Rab13-2, co-immunoprecipitation experiments were performed with Rab13-2 homologs. Pc12 was found to bind selectively to Rab13-3 and Rab13-4, which share 98% and 91% amino acid sequence similarity with Rab13-2, respectively (Figure 3B). This suggests that Pc12 preferentially interacts with specific Rab13-2 homologs. Interestingly, one of the Pc12 homologs demonstrated to induce cell death through its transient expression in *N. benthamiana* previously (Li et al. 2019) also interacts with one of the Rab 13 proteins (Rab 13-4), which localizes with the ER and in the nucleus (Li et al. 2022). GFP-Rab13-2 localizes close to the ER (Figure 3C) and colocalizes with the Golgi apparatus (Figure 3D), suggesting a role for Rab13-2 at the ER-Golgi interface. Rab proteins contain two loops in their globular region that undergo allosteric conformational changes depending on whether the Rab protein is bound to GTP or GDP. In the GTP-bound state, these loops move closer together, resulting in a compact structure. In contrast, Rab proteins bound to GDP have a more relaxed structure (Gray et al. 2020). To determine the specific state of Rab13-2 targeted by Pc12, we generated mutants representing the constitutively active (Rab 13-2^Q74L^) or inactive (Rab13-2^S29N^) form of Rab13-2 based on the consensus sequences (Langemeyer et al., 2014; Bourne et al., 1991). In vitro GTPase assays confirmed the predicted enzyme activities of these mutants (Figure 4A). Co-IP assays showed that Pc12 interacts exclusively with the active form of Rab13-2 (Figure 4B). The direct interaction of Pc12 and Rab13-2 was further confirmed by a yeast two-hybrid (Y2H) assay (Figure 4C). In support, no interaction is observed between Rab 13-2 and Pc12ΔC5, highlighting the importance of the C-terminus of Pc12 in facilitating the interaction between Pc12 and Rab13-2.

**Figure 3.**
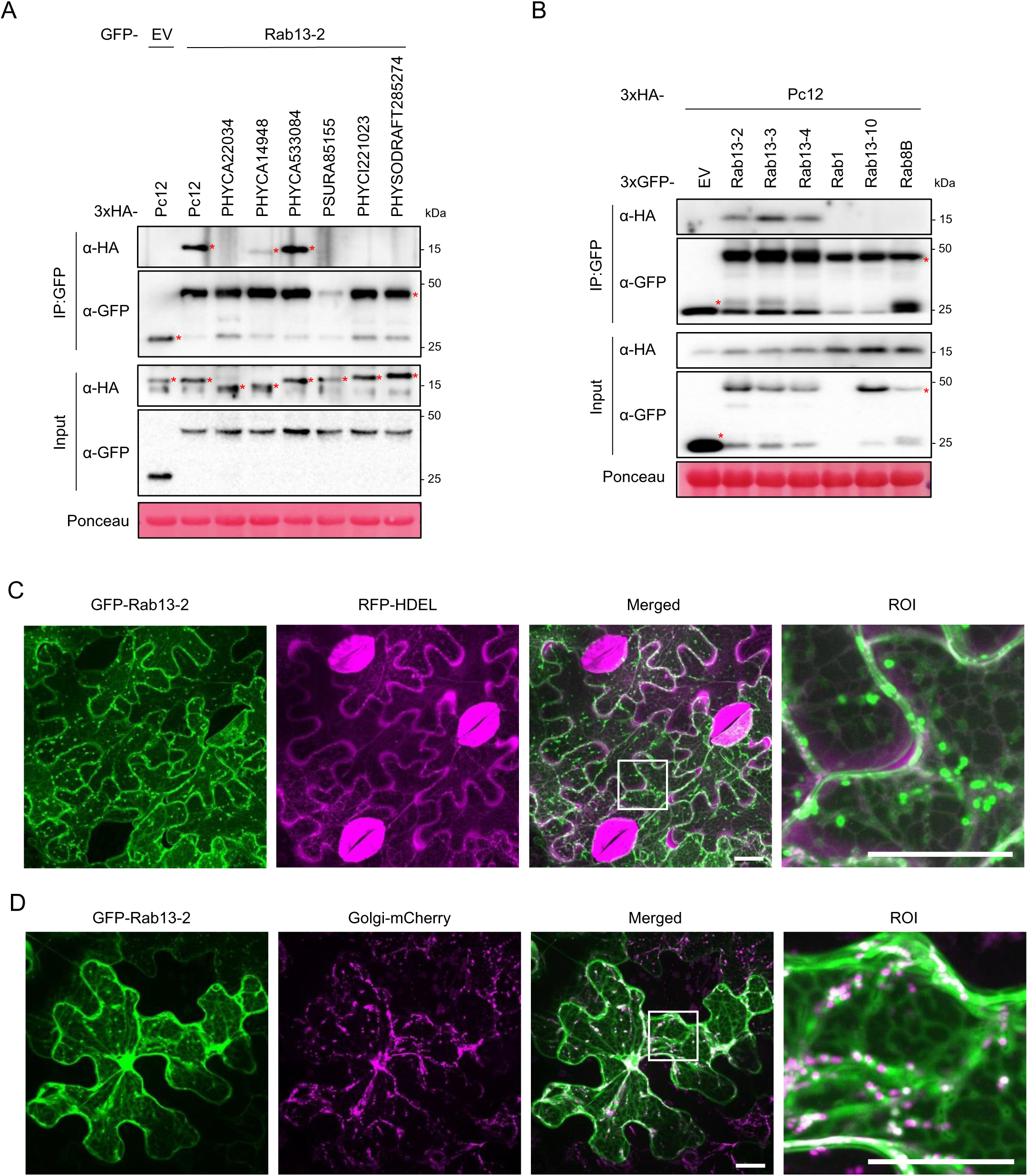
The Pc12 family interacts with the small GTPase Rab13-2 homologs, directly binding to its active form. (A) Rab13-2 interacts with Pc12 homologs, inducing cell death. Rab13-2, with a N-terminal GFP, and 3xHA-Pc12 were transiently expressed in plants. Leaf samples were sampled at 30 hpi. Total protein extracts were subjected to co-IP using anti GFP agarose beads. (3 experimental repeats) (B) Pc12 interacts with the subclade including Rab13-2, -3, and -4. The plants were transiently expressing 3xHA-Pc12 and GFP-Rab13 homologs and sampled at 30 hpi. (3 experimental repeats) (C) GFP-Rab13-2 proteins partially co-localize at the proximity of ER tubules. GFP-Rab13-2 and RFP-HDEL were transiently expressed in the epidermal cells of *N. benthamiana* leaves. GFP was pseudo-colored to green, while RFP displayed in magenta. More than 12 images were acquired from 3 independent experiments. Scale bars, 20 *µ*m. (D) Puncta localization of GFP-Rab13-2 were colocalized with Golgi marker when transiently expressed them in the epidermal cells of *N. benthamiana* leaves. GFP was pseudo-colored to green, while mCherry displayed in magenta. More than 12 images were acquired from 3 independent experiments. Scale bars, 20 *µ*m.

**Figure 4.**
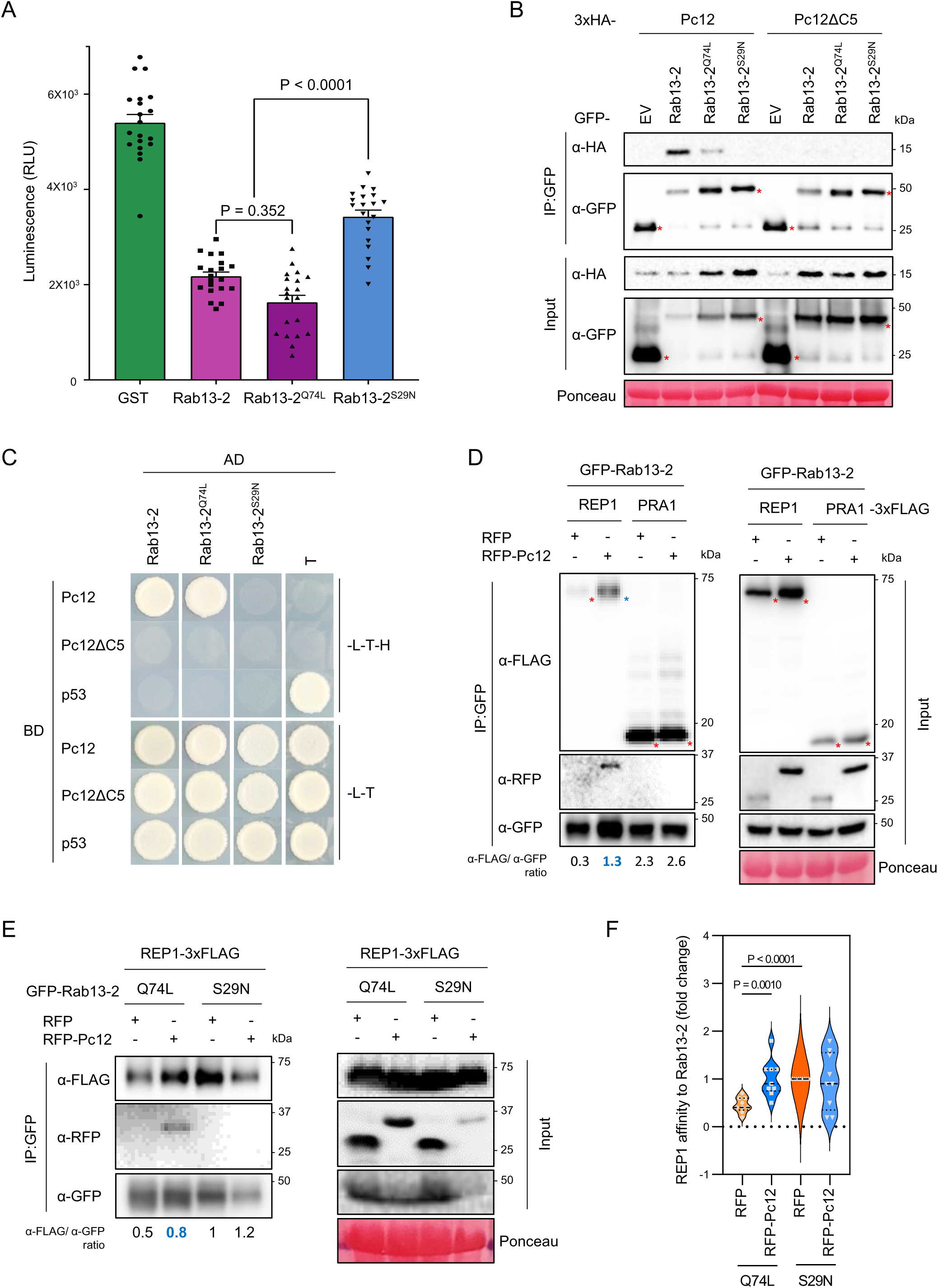
Pc12 directly binds to the active form of Rab13-2, enhancing the interaction between the active form of Rab13-2 and REP1. (A) Intrinsic GTPase activity for Rab13-2 and its mutants Rab13-2^Q74L^ and Rab13-2^S29N^ was measured by using recombinant proteins expressed in *E. coli.* As expected, Rab13-2^Q74L^ showed stronger activity statistically significant, while Rab13-2^S29N^ lost substantial level of activity. Experiments were repeated more than five times with the similar results of the same conclusion. One representative graph is shown with 20 technical replicates. Dots on the graph indicate individual technical replicate. (B) Interaction of Pc12 with Rab13-2 and Rab13-2^Q74L^, not Rab13-2^S29N^ *in planta*. Pc12 and Pc12ΔC5 were transiently co-expressed with Rab13-2, Rab13-2^Q74L^, and Rab13-2^S29N^ in *N. benthamiana*. (C) Physical interaction of Pc12 with Rab13-2 and Rab13-2^Q74L^, not Rab13-2^S29N^ *in vivo*. Yeast cells transformed with GAL4BD-Pc12 or Pc12ΔC5 and GAL4AD-Rab13-2 or mutants were grown on synthetic media lacking LTH or LT for 7 days. The combination of GAL4BD-p53 and GAL4AD-T was used as a positive control. (D) Co-IP assays showing that Pc12 increased binding affinity between Rab13-2 and REP. The plants transiently expressing GFP-Rab13-2, REP- or PRA1-3xFLAG, and RFP or RFP-Pc12 were sampled at 12 hours after a 5% ethanol treatment at 1 day-post-agroinfiltration. Total protein extracts were subjected to co-IP using anti GFP agarose beads. The precipitated proteins were immunoblotted. The level of binding was calculated as the ratio between anti-FLAG and anti-FLAG of IP by imageJ. Red asterisks indicate expected band sizes. (E) Pc12 strengthen the weak interaction between REP and Rab13-2^Q74L^. The plants were infiltrated with an agrobacterium carrying REP-3xFLAG and an agrobacterium containing two plasmids, one plasmid with RFP or RFP-Pc12 and the other with GFP-Rab13-2^Q74L^ or Rab13-2^S29N^, in a 1:1 ratio in *N. benthamiana*. Samples treated with 5% ethanol at 1 dpi were collected at 12 hours post treatment. The level of binding was calculated as the ratio between anti-FLAG and anti-FLAG of IP by imageJ. Red asterisks indicate expected RFP and RFP-Pc12 band sizes. (F) Statistical graph depicting the results of 8 repeated experiments for (E).

### The complex of Pc12 and Rab13-2 mimics the inactive conformation of Rab13-2

Rab proteins are involved in vesicle budding, trafficking, and fusion processes by interacting with or recruiting other proteins (Stenmark 2009; Pylypenko et al. 2018; Nielsen et al. 2020). In the case of Rab13-2, potential interactors were identified using the STRING database (Supplemental Table 3). Among the interactors, the Rab escort protein 1 (REP1) and the prenylated Rab acceptor 1 (PRA1) were found to interact with Rab13-2 by co-IP screening (Supplemental Figure 4). REP1 has a dual role: it acts as a chaperone to facilitate the prenylation process of Rab and as a carrier to deliver Rab to its target membrane (Guo et al. 2008). PRA1 acts as a receptor for prenylated Rab proteins, promoting their binding to membranes (Figueroa et al. 2001). We hypothesized that Pc12 might affect the interaction between Rab13-2 and its interactors, REP1 or PRA1. To confirm the effect of Pc12 on these interactions, we performed an immunoprecipitation experiment. Rab13-2 and its interactors were continuously expressed in a plant, while Pc12 was induced by ethanol treatment at 1 dpi (day post-infiltration) to ensure that the action of Pc12 did not affect the accumulation of newly synthesized proteins. The results showed that Pc12 did not form a complex with Rab13-2 in the presence of PRA1 (Figure 4D), suggesting that Pc12 does not interact with Rab13-2 associated with PRA1. On the other hand, Pc12 interacts with Rab13-2 associated with REP1 (Figure 4D). Furthermore, the interaction between Rab13-2 and REP1 was enhanced by the expression of Pc12 (blue asterisk, α-FLAG/α-GFP = 1.3, Figure 4D). This suggests that Pc12 plays a regulatory role in the association between Rab13-2 and REP, potentially affecting their cellular functions and signaling pathways. Pc12 displays a preference for binding to the active form of Rab13-2 (Rab 13-2^Q74L^, Figure 4B, C). However, REP1 has been shown to interact predominantly with the inactive form (Stenmark 2009, Supplemental Figure 5). Therefore, we hypothesized that binding of Pc12 to the active form of Rab13-2 might alter the structural conformation similar to the inactive state of Rab13-2, thereby facilitating the recruitment of REP1 to active Rab13-2. To validate this hypothesis, we performed a co-IP with Pc12, Rab13-2^Q74L^, Rab13-2^S29N^, and REP1. Remarkably, REP1 showed a significant association with Rab13-2^Q74L^ in the presence of Pc12, compared to the control RFP (Figure 4E). This experiment was repeated eight times, all with consistent results (Figure 4F, Supplemental Figure 6). This finding indicates that Pc12 binds to the active state of Rab13-2, thereby facilitating the recruitment of REP1. This recruitment may alter the dynamics of Rab13-2 GTPase activity, ultimately leading to the disruption of vesicle trafficking.

### Pc12 interaction with Rab13-2 alters ER-Golgi trafficking

Rab proteins are known for their distinct localization within the plant endomembrane network (Stenmark 2009; Nielsen et al. 2020). The hypervariable C-terminal region of Rab proteins contributes to the specificity of membrane delivery for certain Rab proteins (Figueroa et al. 2001; Pylypenko et al. 2018). Previously, a Pc12 isoform was identified that interacts with a Rab 13 subfamily isoform localizes to the ER (Li et al. 2022). To gain insight into the localization of Pc12 within host cells, we transiently expressed EGFP-Pc12 and EGFP-Pc12ΔC5 (without the signal peptide and RXLR-EER motifs) under the control of an ethanol-inducible promoter. Due to the rapid induction of cell death by Pc12, cytosolic Pc12 expression was observed in the early time point following 1% ethanol spray treatment to determine its subcellular localization (Figure 5A). Interestingly, in contrast to the previously studied isoform (Li et al. 2022), the Pc12 proteins are defused in the cytoplasm with preferential accumulation closer to the ER and Golgi (Figure 5A). We attempted to confirm the Pc12 localization in the vicinity of the ER and Golgi apparatus by co-expressing ER marker (Chakrabarty et al. 2007) and Golgi marker (Nelson et al. 2007, Park et al. 2017). Surprisingly, we observed that cells showed a peculiar punctate localization of RFP-HDEL in the cell (Figure 5B), whereas the reticular morphology of the ER visualized by RFP-HDEL is unchanged under the expression of EGFP or EGFP-Pc12ΔC5 (Figure 5B). In addition, the cis-Golgi marker, soybean α-1,2 mannosidase I fused to mCherry (Nelson et al. 2007), was trapped in the ER under Pc12 expression (Figure 5C). The mislocalization of these two markers under the Pc12 expression suggests that perturbation of the Rab13-2 dynamics by Pc12 interaction strongly affects vesicle trafficking from the ER to the Golgi.

**Figure 5.**
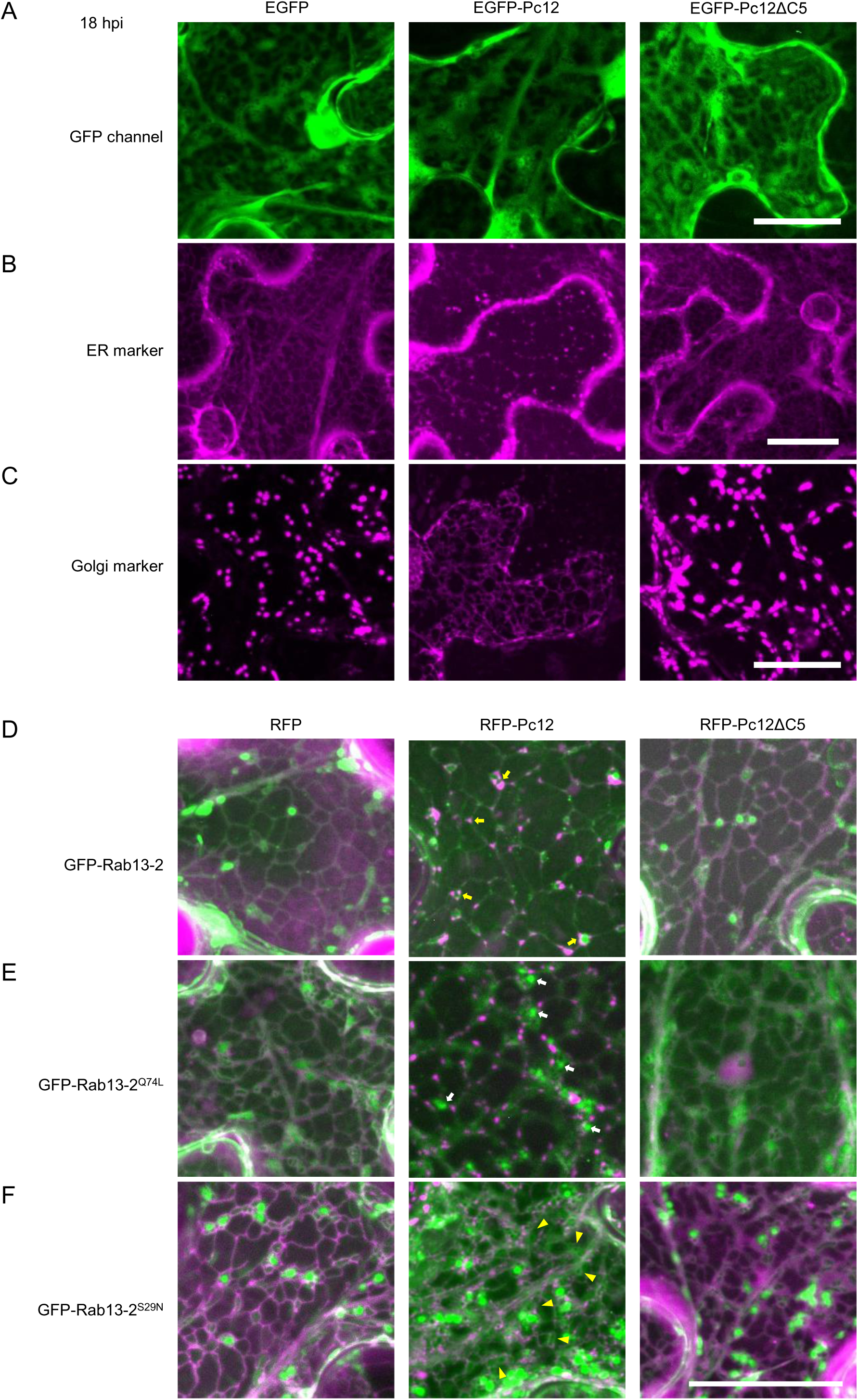
Pc12 interrupts trafficking of RFP-HDEL and *Cis-*Golgi marker and relocates Rab13-2, mimicking the inactive state of Rab13-2. (A) Pc12 and Pc12ΔC5 are localized in the cytoplasm. GFP-Pc12 as well as GFP-Pc12ΔC5, a control GFP were infiltrated in the leaves of a transgenic *N. benthamiana* expressing RFP-HDEL as an ER marker (Magenta). Transient expressions of those proteins were imaged by a spinning disc confocal microscope. More than 60 z-images in 0.3 *µ*m step size were superimposed by a maximum z-projection function in ImageJ. The brightness of the original micrograms was enhanced to improve the visibility of subcellular compartments. This processing does not change the conclusion drawn from the images. GFP was pseudo-colored to green, while RFP displayed in magenta. More than 12 images were acquired from 3 independent experiments. Scale bars, 20 *µ*m. (B) Discontinuity and puncta localization of RFP-HDEL under the expression of Pc12. GFP, GFP-Pc12 and GFP-Pc12ΔC5 were transiently expressed in transgenic *N. benthamiana* expressing RFP-HDEL. More than 60 z-images in 0.3 *µ*m step size were acquired by a spinning disc confocal microscope. Images were superimposed by a maximum z-projection function in ImageJ. The brightness of the original micrograms was enhanced to improve the visibility of subcellular compartments. This processing does not change the conclusion drawn from the images. RFP displayed in magenta. More than 12 images were acquired from 3 independent experiments. Scale bars, 20 *µ*m. (C) Rapid trafficking of *cis*-Golgi marker, Soybean α-1,2 mannosidase I fused to mCherry, was interrupted by Pc12 expression. GFP, GFP-Pc12 and GFP-Pc12ΔC5 were transiently expressed with Golgi marker *N. benthamiana*. Image acquisition and processing were identical to those for (B). The brightness of the original micrograms was enhanced to improve the visibility of subcellular compartments. This processing does not change the conclusion drawn from the images. mCherry displayed in magenta. More than 12 images were acquired from 3 independent experiments. Scale bars, 20 *µ*m. (D-F) For more accurate localization and dynamics of Rab13-2 and its mutants, transgenic *N. benthamiana* plants expressing both RFP-HDEL and GFP-Rab13-2 (D), GFP-Rab13-2^Q74L^ (E), or GFP-Rab13-2^S29N^ (F) were generated. Stable expressions of those proteins in the leaf epidermal cells were imaged by a spinning disc confocal microscope. More than 60 z-images in 0.3 *µ*m step size were superimposed by a maximum z-projection function in ImageJ. The brightness of the original micrograms was enhanced to improve the visibility of subcellular compartments. This processing does not change the conclusion drawn from the images. GFP was pseudo-colored to green, while RFP displayed in magenta. More than 12 images were acquired from 3 independent experiments. Scale bars, 20 *µ*m.

To better understand Rab13-2 function and Pc12 interference with vesicle trafficking at the ER-Golgi interface, we generated transgenic *N. benthamiana* plants expressing Rab13-2 together with the ER marker. As we observed with transient expression of GFP-Rab13-2 (Figure 3C, D), Rab13-2 proteins colocalized in the vicinity of ER and Golgi puncta structures (Figure 5D, top panel). Interestingly, under Pc12 expression, puncta structures shown by ER markers were located at ER tubules and often accumulated near Golgi (Figure 5D, yellow arrows). ER marker consist of mCherry, a red fluorescent protein fused to a topic signal peptide at the N-terminus and an ER retention signal peptide (HDEL) at the C-terminus (Park et al. 2017). RFP-HDEL typically displays a reticular ER morphology when observed under a microscope. This is because newly synthesized proteins containing HDEL are first synthesized within the ER lumen and then transported to the Golgi apparatus. They then return to the ER via signal peptide recognition (Alvim et al. 2023). The punctate accumulation of RFP-HDEL at the ER tubules and the proximity to the Golgi induced by Pc12 suggest that it may be caused by interference with the mCherry proteins secreted from the ER to the Golgi.

Rab small GTPases continuously switch enzyme properties between their GTP-bound and GDP-bound forms. Conventionally, the active state of Rab proteins was thought to be localized to membranes, while the inactive state was predominantly found in the cytoplasm, as dictated by their respective functions (Stenmark, 2009; Nielsen, 2020). To validate the localization of Rab13-2 mutants, we generated transgenic *N. benthamiana* expressing the constitutive active form Rab13-2^Q74L^ or the constitutive inactive form Rab13-2^S29N^ together with an ER marker (Figure 5E and F, respectively, supplemental Figure 7). Notably, the constitutive inactive form Rab13-2^S29N^ was frequently observed at the Golgi apparatus (Figure 5F, supplemental Figure 7C), but the constitutive active form Rab13-2^Q74L^ did not show the Golgi localization (Figure 5E, supplemental Figure 7B), contradicting the general expectation of Rab small GTPase dynamics of membrane binding upon their activation by GTP recruitment (Stenmark, 2009; Nielsen, 2020). The localization pattern of Rab13-2^Q74L^ is more similar to that of RabE1d^Q74L^ and Rab8^Q74L^ (Speth et al., 2009; Pandy et al., 2021). Since Rab proteins are initially synthesized in a structurally stable active state, the host interactors responsible for Rab prenylation and membrane anchoring are unable to bind to the mutant due to its abnormal conformation.

With this in mind, we investigated whether the localization of Rab13-2 might be altered upon coexpression of Pc12. We observed that larger punctate localizations of GFP-Rab 13-2^Q74L^ were frequently observed upon expression of Pc12 (white arrows, Figure 5E), whereas they were not observed with RFP or Pc12ΔC5. Furthermore, a more diffuse localization of GFP-Rab13-2^S29N^ in the vicinity of the ER was observed under the Pc12 expression (yellow arrowheads, Figure 5F). The results suggest that Pc12 alters the properties of Rab13-2^Q74L^ to resemble those of Rab13-2^S29N^ and effectively recruiting REP1 and consequently localizing it to the Golgi apparatus.

### Pc12 interference with ER-Golgi vesicle trafficking primarily affects the secretory pathway and induces ER stress leading to necrotic cell death

The altered localization of Rab 13-2^Q74L^ with Pc12 indicates that the rapid dissociation of active Rab proteins and their return to the ER periphery may be disrupted under Pc12 expression. Displacement of ER marker and Golgi marker under Pc12 expression (Figure 5B and C) also suggested that the alteration of Rab13-2 dynamics might disrupt trafficking from the ER to the Golgi. Therefore, we observed the localization of two subcellular compartment markers (Figure 6A and B). Notably, an apoplast marker of moxVenus fluorescent proteins, fused to an apoplastic signal peptide at the N-terminus to secrete the fluorescent protein outside the plasma membrane (ApoSP-moxVenus, Balleza et al. 2017), is trapped in the ER in the presence of RFP-Pc12 (Figure 6A). This observation suggests that Pc12 disrupts vesicle trafficking at the ER-Golgi interface, leading to changes in the localization of the markers.

**Figure 6.**
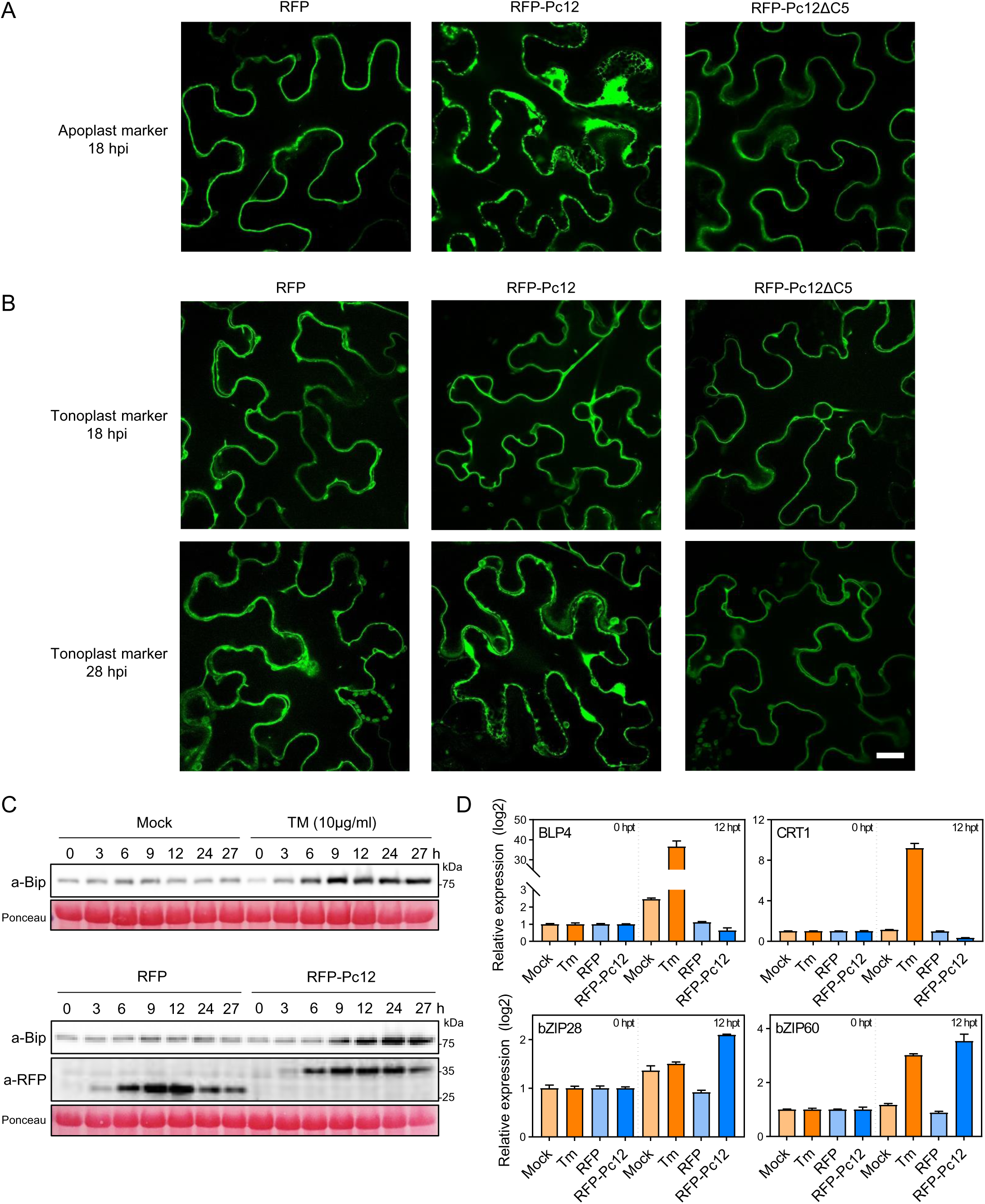
Pc12 inhibits the secretion from ER to Golgi, triggering ER stress-mediated cell death. (A) Apoplastic membrane targeted moxVenus (apoSP-moxVenus-TM) remained in ER in the existence of Pc12. RFP-Pc12, RFP-Pc12ΔC5, and RFP driven by the ethanol promoter were induced by 1% ethanol treatment. Pc12 expression inhibited the secretion of the marker proteins resulted in their accumulation in the ER. Z-images for thicker than 3 *µ*m sections in 0.75 *µ*m step size were acquired by a laser scanning confocal microscope. Images of the corresponding sections were processed to improve the brightness for the clarity. This processing does not change the conclusion drawn from the images. moxVenus was pseudo-colored to green. More than 12 images were acquired from 3 independent experiments. Scale bars, 20 *µ*m. (B) Tonoplast marker (*r*TIP, vacuolar membrane protein, fusion at the N-terminal of moxVenus) was transiently expressed in leaf epidermis of *N. benthamiana*. Trafficking of tonoplast marker takes longer time than other proteins. Thus, tonoplast marker was infiltrated 2 days ahead than RFP, RFP-Pc12, or RFP-Pc12ΔC5. Images were acquired 18 hours (top panel), 28 hours (bottom) panels, and 36 hours (supplemental figures 8B) after spraying 1% ethanol. Images were process with the same procedure described in (D). moxVenus was pseudo-colored to green. More than 12 images were acquired from 3 independent experiments. Scale bars, 20 *µ*m. (C) Accumulation of the ER stress marker protein (Bip) in response to Pc12 expression. Plants were treated with mock and Tm (10 ug/ml) and sampled over time. Plants expressing RFP and RFP-Pc12 after 5% ethanol treatment were sampled over time. (D) Transcripts level of UPR-related transcription factor and ER chaperon genes upon expression of Pc12. In (C), total RNA was extracted at 0 and 12 hours after tunicamycin or ethanol treatment. UPR-related genes, ER chaperon genes (*BLP4* and *CRT1*) and transcription factor genes (*bZIP28* and *bZIP60*), were normalized with *NbEF1a* in qRT-PCR.

Interestingly, the localization of a tonoplast marker, moxVenus fused to γ-TIP (Nelson et al. 2007), was not significantly affected approximately 18 h after induction of Pc12 expression (Figure 6B, top panel), while the apoplast marker trafficking was completely blocked in the ER (Figure 6A, Supplemental Figure 8A). At a later time point, 28 hours after Pc12 induction, dead cells are often observed under the microscope, and the tonoplast markers are observed in the ER (Figure 6B, lower panel). Later, around 36 h after Pc12 induction, most of cells are already completely dead, but tonoplast markers rarely maintain their integrity (Supplemental Figure 8B). An alternative pathway of vesicle trafficking from the ER to the vacuole, bypassing the Golgi, has been investigated as an important route for ER-vacuole trafficking (Rojas-Pierce, 2013). This observation suggests that the primary function of Rab13-2 in the secretory pathway is through the ER-Golgi pathway. Thus, perturbation of this pathway by Pc12 alternation of Rab13-2 dynamics could induce a significant traffic jam of secretion at the ER-Golgi interface, ultimately inducing ER stress-mediated necrotic cell death. Interestingly, the mutualistic fungus *Piriformospora indica* has been shown to induce ER stress-induced cell death when it colonizes *Arabidopsis* roots (Qiang et al. 2012). Furthermore, the altered ER morphology induced by *P. indica* infection is similar to that induced by Pc12 expression (Qiang et al. 2012, Figure 5B). Therefore, we compared the responses induced by Pc12 and tunicamycin, a known inducer of ER stress. These responses include the accumulation of ER chaperone-binding immunoglobulin protein (BiP) and the transcription of unfolded protein response (UPR) genes (Cao and Kaufman 2012; Angelos et al. 2017; Adams et al. 2019; Yang et al. 2014). Following application of 5% ethanol to the leaves, ethanol-induced expression of RFP-Pc12 and RFP resulted in the accumulation of both proteins over time. As hypothesized, RFP-Pc12 induced a gradual accumulation of BiP over time, similar to the effects of tunicamycin treatment (Figure 6C). To assess the expression of UPR-related genes, samples were taken at 0 and 12 hours post-treatment. Tunicamycin treatment resulted in the upregulation of transcripts for both ER chaperones (BiP-like protein *BLP4* and calreticulin *CRT1*) and ER stress-related transcription factors (*bZIP28* and *bZIP60*). However, Pc12 expression did not show a similar pattern of upregulation of ER chaperone transcripts. Instead, Pc12 induced an increase in the expression of transcription factors (Figure 6D). These results suggest that Pc12 induces a distinct form of ER stress that is different from the effects of tunicamycin.

### A specific residue of Rab13-2 is crucial for its interaction with Pc12 and its mutation evading targeting by Pc12 affect to the virulence of *P. capsici*

The failure of effectors to suppress plant immunity by manipulating the function of host targets is thought to contribute factor to non-host resistance (NHR) (McLellan et al. 2022). Thus, breaking down the interaction between Pc12 and Rab13-2 could reduce the susceptibility of plants to pathogens. To obtain a better estimate of the biochemical nature of the interaction between Pc12 and Rab13-2, we used the AlphaFold2 program (Jumper et al. 2021) to identify a pivotal residue on Rab13-2 that interacts with the C-terminal AHQMG residues of Pc12. Rab13-2 was predicted to have a globular structure, whereas Pc12 displayed a stack of three parallel α-helices connected by a short helical linker (Figure 7A). A docking model proposed that the three parallel α-helices of Pc12 interact with the globular region of Rab13-2 (Figure 7A). In particular, L90, A91, and M94 at the C-terminus of Pc12 were predicted to engage with threonine at the 47th position of Rab13-2 (Figure 7B). Remarkably, replacement of threonine with alanine (Rab13-2^T47A^) abolished the Rab13-2 interaction with Pc12 (Figure 7C) without affecting its ability to bind REP1 and PRA1 (Figure 7D-E).

**Figure 7.**
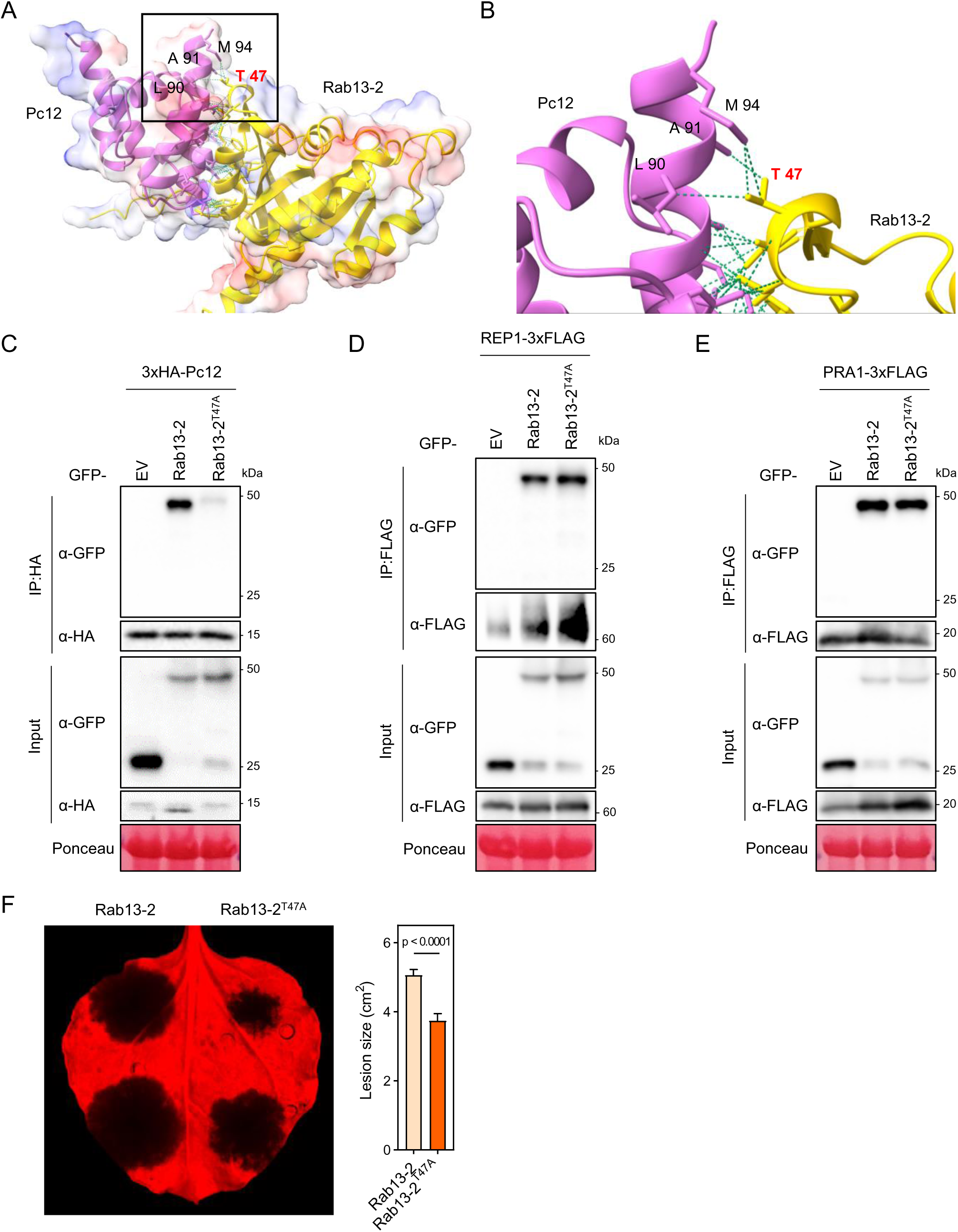
A mutation in a key Rab13-2 residue weakens Pc12 binding without affecting REP1 and PRA1 binding, compromising *P. capsici* virulence. (A-B) The predicted structures of Pc12 and Rab13-2 interaction by AlphaFold2. (B) zooms in on the black box in (A). The green lines depict the intracellular bonds between Pc12 and Rab13-2. (C -E) *In planta*, a co-immunoprecipitation assay with Rab13-2, Rab13-2^T47A^, and Pc12 (REP1 or PRA1). Total protein extracts were precipitated with anti HA magnetic beads or anti FLAG agarose. The precipitated protein and total protein were detected by western blotting. (F) *P. capsici* inoculation on the leaves expressing GFP-Rab13-2 and GFP-Rab13-2^T47A^. Agrobacterium carrying GFP-Rab13-2 and GFP-Rab13-2^T47A^ was infiltrated in *N. benthamiana*, followed by *P. capsici* inoculation at 1 dpi. Images were taken at 72 hpi, and the lesion size was measured by ImageJ (N = 31 - 32, 3 repeats).

These results are important in considering the potential of Rab proteins as a target for engineering plants resistant to *P. capsici*. To evaluate the applicability of the Rab13-2^T47A^ mutation for resistance to *P. capsici*, *N. benthamiana* plants expressing Rab13-2^T47A^ were inoculated with *P. capsici*. At 72 hpi, as the disease progressed, lesion size was significantly reduced in Rab13-2^T47A^-expressing leaf compared to Rab13-2-expressing leaf (Figure 7F). Considering that the effect of endogenous Rab13-2 might diminish the effect of the expression of Rab13-2^T47A^ expression, this result is significant to suggest that Pc12 is a key player in determining the necrotrophic phase and Rab13-2^T47A^ may serve as a contributing factor to delay the onset of the necrotrophic disease progression.

## Discussion

### Pc12 may serve as a transition effector, orchestrating the transition from the biotrophic to the necrotrophic life cycle of *P. capsici*

*P. capsici* is classified as a hemibiotrophic pathogen, characterized by a biphasic life cycle. It begins with a biotrophic phase in which it strategically establishes itself within the host plant. It then transitions to a necrotrophic phase aimed at extracting nutrients from dead cells and releasing spores for subsequent dissemination. (Kamoun et al. 2015; Parada-Rojas et al. 2021; Quesada-Ocampo et al. 2023). Although researchers have uncovered the role of RXLR effectors in increasing susceptibility by interfering with host cell physiology or triggering defense responses through interactions with R proteins during the biotrophic phase, our understanding of how these effectors induce necrosis in host cells remains quite limited. Recently, a case of necrosis was reported in which a *P. infestans* RXLR effector induced nucleolar inflammation and interfered with pre-rRNA 25S processing, leading to necrosis through disruption of protein translation (Lee et al. 2023). This study identified the RXLR effector Pc12 as a critical factor in necrosis, which causes severe ER stress by preventing Rab13-2 recycling in vesicle trafficking. Furthermore, bypassing Pc12 targeting by Rab13-2^T47A^ led to a reduction in necrotrophic lesion size, highlighting that Pc12-induced necrosis contributes to the necrotrophic life cycle of the pathogen. Interestingly, Pc12 expression was highly upregulated at 6 hpi, while the transcripts of NPP1, a necrotrophic marker, gradually increased at 12 hpi (Supplemental Figure 1). This raises the question of why Pc12 transcription increased early in infection despite its role in inducing necrosis. In the experiment using the ethanol-inducible promoter, the accumulation of the Pc12 protein was detectable by immunoblots from 3 h after 5 % ethanol treatment (Figure 6C), and cell death was observed by microscopy before 24 h after 1 % ethanol treatment by microscopy (Supplemental Figure 8). Given the time delay between gene transcription and necrosis induction, it could be inferred that Pc12-induced necrosis serves as an initiator for entry into the necrotrophic phase.

### The emergence of Pc12 may benefit *P. capsici* by broadening its host range and promoting survival

Within the genus *Phytophthora*, only *P. capsici* possesses multiple copies of at least 8 homologs of Pc12 with the potential to induce necrosis. In contrast, *P. ramorum*, *P. cinnamoni*, and *P. sojae* lack this specificity (Supplemental Table 1, Supplemental Figure 2). *P. infestans* lacks a Pc12 homolog, despite its close evolutionary relationship with *P. capsici* (Lamour et al., 2012; Sharma et al., 2015). This suggests that the Pc12 family likely evolved after the emergence of *P. capsici* and underwent lineage-specific expansion within the species. Given the distinctive characteristics of *P. capsici*, the emergence of Pc12 has led to two positive outcomes. First, Pc12 disrupts vesicle trafficking in the basal plant physiology, allowing *P. capsici* to infect a wider range of hosts compared to *P. infestans* and *P. sojae*. Second, Pc12 rapidly induces necrosis, causing *P. capsici* to shorten its biotrophic phase to approximately 30 hpi, in contrast to *P. infestans* and *P. sojae* (2 days). This acceleration could help *P. capsici* to colonize well before the onset of the host defense response, allowing the pathogen to evade early immune responses. With these advantages conferred by Pc12, *P. capsici* may have the potential to cause extensive damage to a wide range of hosts, often resulting in crop losses of up to 100%.

### Pc12 disrupts the Rab13-2 cycle by increasing the affinity of REP1 to the active structure of Rab13-2

Numerous studies have reported that pathogenic effectors target vesicle-associated proteins to disrupt the secretion of defense-related proteins or to subvert autophagy to obtain nutrients for pathogen nutrition (Gu et al. 2017; Pandey et al. 2021; Petre et al. 2021; Yuen et al. 2023). In our research, we have shown that Pc12 targets Rab13-2 and disrupts vesicle trafficking between the ER and the Golgi. This disruption leads to the accumulation of apoplastic proteins in the ER lumen (Figure 6A, Supplemental Figure 8A). Similarly, the viral replicase p27 of *Red clover necrotic mosaic virus* (RCNMV) inhibits vesicle trafficking between the ER and Golgi by translocating Arf1 for COPI and Sar1 for COPII to the ER, resulting in the accumulation of secretory proteins in the ER (Hyodo et al. 2013; Hyodo et al. 2014). We have also discovered a novel strategy for the RXLR effector Pc12. It mimics the inactive conformation of Rab13-2 by binding to the active structure of Rab13-2, thereby increasing the affinity of REP1 for the active form of Rab13-2 (Figure 4, 8). This strategy is similar to that of brefeldin A, a fungal toxin that increases the affinity of GEFs to bind to GDP-bound Arf1, ultimately disrupting the cycling of Arf1 between GDP and GTP (Niu et al. 2005). Given the shared goals of a viral protein and a fungal toxin, Pc12 could contribute significatnly to the pathogenicity of the pathogen.

### Necrosis caused by Pc12 results from a specific form of severe ER stress characterized by hyper-accumulation of proteins

Both animals and plants have evolutionarily conserved ER stress responses to a variety of stimuli (Oakes et al. 2015; Angelos et al. 2017). When ER dysfunction is excessive or prolonged, cells initiate death signaling, similar to a survival strategy in multicellular organisms, aimed at eliminating dysfunctional cells (Oakes et al. 2015; Simoni et al. 2022). ER stress-induced cell death has been extensively studied in animals, particularly in the context of diseases such as Alzheimer’s and Parkinson’s (Oakes et al. 2015). However, our understanding of ER stress-induced cell death in plants is still evolving and some aspects have yet to be fully elucidated. Pc12-induced inhibition of Rab13-2 recycling in vesicle trafficking leads to severe ER stress, ultimately culminating in necrosis. According to Yang et al. 2014, the transcription factor NAC089, transcribed by bZIP28/60, plays a key role in controlling ERstress-induced programmed cell death in plants by triggering the expression of PCD-related genes (NAC094, MC5, and BAG6) (Yang et al. 2014). Notably, Pc12 induces a substantial upregulation of bZIP28/60, but does not lead to a corresponding increase in the expression of bZIP28/60-downsteam pathway genes (UPR-related genes, Figure 6C) and PCD-related genes (not data shown). Coupled with the Pc12-induced disruption of vesicle trafficking between the ER and the Golgi, this implies that Pc12 prevents bZIP28 translocation to the Golgi and prevents bZIP60 splicing in the absence of signaling of unresolved ER stress. This results in the repression of genes regulated by bZIP28/60 (Figure 6C). Furthermore, Ko et al. (2023) recently reported that ABI5 activates the transcription of bZIP60 under ER stress, suggesting the interpretation that Pc12 induces the upregulation of bZIP28/60 independent of bZIP28/60 translocation to the nucleus (Ko et al. 2023). However, the movement of ABI5 under ER stress remains unknown. Taken together, the failure to alleviate ER stress by Pc12 may lead to an excessive accumulation of proteins in the ER, effectively acting as a cellular “time bomb” (Figure 8).

**Figure 8.**
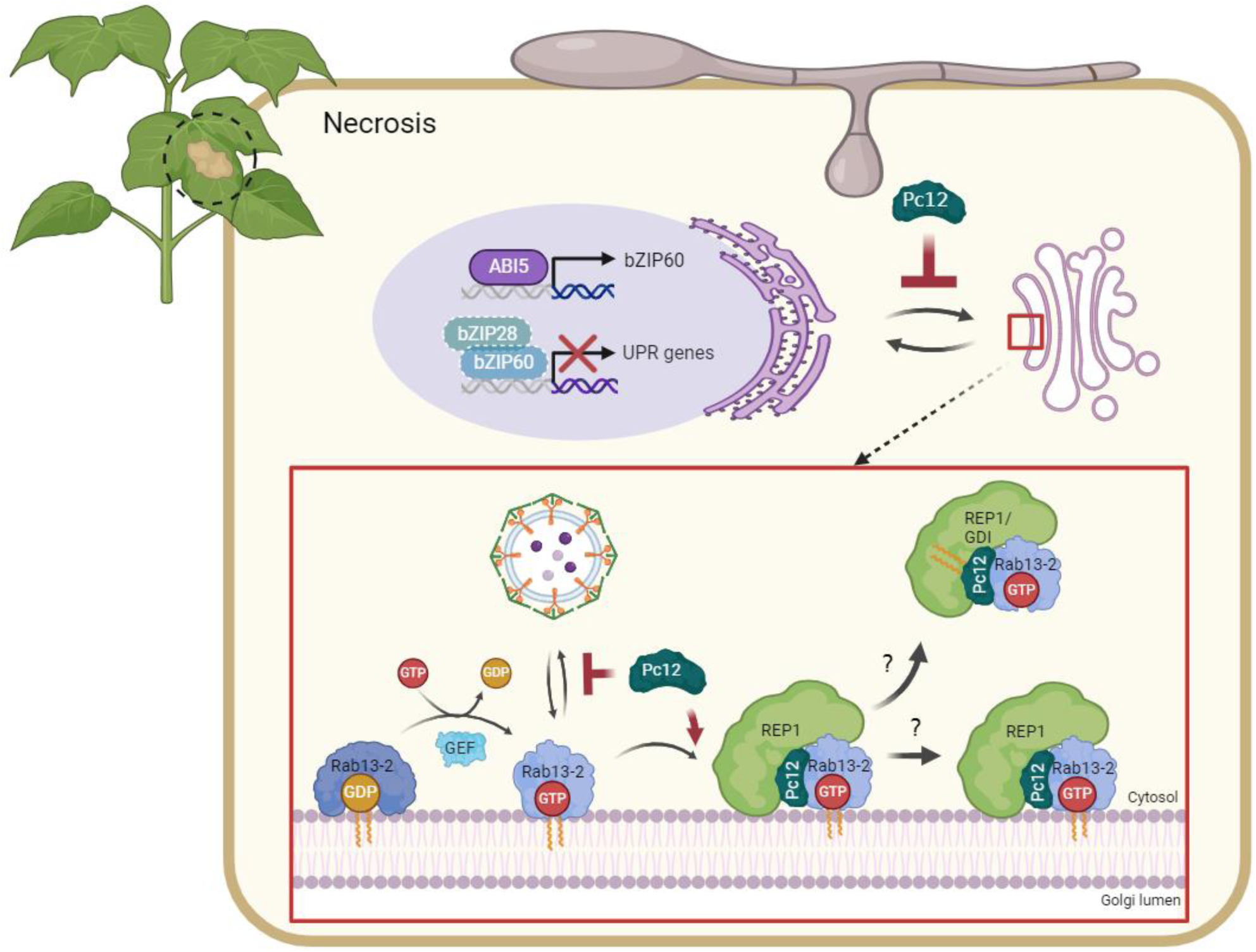
Summary model of the inhibition of Pc12 in Rab13-2-mediated vesicle trafficking. Prenylated Rab13-2 in its GDP-bound state is inserted to a membrane by Rab escort protein 1 (REP1). The guanine nucleotide exchange factor (GEF) exchanges GDP for GTP on Rab13-2 to initiate vesicle formation for vesicle trafficking. Pc12, secreted from *P. capsici*, binds to GTP-bound Rab13-2, subsequently recruiting REP1 to facilitate the extraction of the complex from the membrane or its retention on the membrane, consequently impeding vesicle formation mediated by Rab13-2. Figure was created using BioRender.com.

### The Rab13-2 point mutation has the potential to manipulate plant resistance to *P. capsici*

The underlying molecular mechanism of non-host resistance is thought to arise from 1) pre-existing barriers that prevent pathogen establishment, 2) the recognition of pathogen effector proteins that induce robust immunity, and 3) the inability of pathogen effectors to suppress immunity induced by molecular patterns (McLellan et al. 2022). The inability of effectors to manipulate host targets for pathogen colonization results in a hostile environment, leading to the failure of the pathogen to adapt to the host. Importantly, this implies that the evasion strategy against effectors targets, acting like an orchestra, results in robust resistance, allowing conversion from host to non-host plants. We identified the specific residue in Rab13-2 that avoids Pc12 targeting by AF2. Mutation of this residue reduced the pathogenicity of *P. capsici*. This study provides insight into the idea that host plants can be engineered to resist *P. capsici* through the synergistic effects of different mutations that bypass other effectors that collectively contribute to susceptibility.

## Materials and methods

### Plant materials and growth conditions

*N. benthamiana, N. tabacum* cv. Samsun, and *Solanum lycopersicum* cv. Heinz seeds were directly sown into damp horticultural bed soil (Biogreen, Seoul, Korea) and cultivated within a walk-in chamber at 22-24℃ under a 16h/8h (day/night) cycle. *Capsicum annuum* cv. ECW seeds were sterilized for 1 min in a 0.1% sodium hypochlorite solution and germinated in darkness at 30℃ for 7 days. Following this, the pepper seedlings were transplanted into soil and grown within the same chamber. For virus-induced gene-silencing assays, 3-week-old *N. benthamiana* plants were utilized. Transgenic *N. benthamiana* plants expressing RFP fused to HDEL (ER marker, Chakrabarty et al. 2007) and GFP-Rab13-2, GFP-Rab13-2^Q74L^, and GFP-Rab13-2^S29N^ transgenic plants were purchased from the Nicotiana Genetic Stock Center (NGSC). Plants were grown at 22-24°C for 4-5 weeks prior to the corresponding experiments.

### *P. capsici* culture condition and inoculation assays

*P. capsici* strain 40476 was cultured on V8 agar medium for 7 days at 23℃ in darkness. The mycelia were scraped and then incubated under light for 12 hours. To induce the release of zoospores, sporangia were harvested and placed in distilled water, incubating at 4℃ for 30 minutes, followed by 23℃ for 30 minutes. A total of 500 zoospores were inoculated on the detached *N. benthamiana* leaves through droplets. The leaves were placed at 25℃ for 2 days under deem light. The lesion areas were measured at 2 dpi.

### Constructs and markers preparation

RXLR effector and their mutants were amplified, incorporating an N-terminal 3xHA epitope excluding signal peptide and RXLR-EER motifs. Rab proteins and REP1 or PRA were amplified with N-terminal GFP and 3xFLAG, respectively, employing the overlap PCR method (Kim et al. 2017). GOIs were inserted into pCambia2300-LIC vector using the ligation-independent cloning method (Oh et al. 2010). RFP and RFP-Pc12 were amplified with the attB site to insert into ethanol-inducible expression vector using Gateway cloning (Invitrogen, USA). The primer sequence used for construction are presented in Supplementary Table 3. mCherry targeted to Golgi generated previously was used for Golgi marker (Addgene ID 97401, Park et al. 2017). The membrane bound apoplastic marker constructs were generated by inserting a signal peptide of *At*Chitinase in front of moxVenus yellow fluorescence protein sequences (Baleza et al. 2017, Berthold et al. 2019). A vacuolar membrane maker, TONO-moxVenus was generated by *r*TIP, vacuolar membrane protein, fusion at the N-terminal of moxVenus (Nelson et al. 2007). Plant expression constructs of Rab variants, apoplastic side membrane marker, and Tonoplast marker were deposited to the Addgene (Rab13-2, ID:213473; Rab 13-2^Q74L^, ID:213474; Rab13-2^S29N^, ID:213475; APO-moxVenus, ID: 220490, TONO-moxVenus, ID: 220491) constructs are deposited in Addgene.

### Agroinfiltration and quantification of cell death assays

*Agrobacterium tumefaciens* GV3101 strain or GV2260 containing the various constructs were cultivated overnight at 28℃ in LB medium supplemented with appropriate antibiotics. The cells were harvested through centrifugation and then resuspended in an infiltration buffer (10 mM MgCl_2_, 10 mM MES (pH 5.6), and 200 μM Acetosyringone). Resuspensions were adjusted to an optical density (OD600) of 0.1-0.3 and leaves of 4-week-plants were agroinfiltrated for transient expression. To quantify the degree of cell death, *N. benthamiana* leaves were detached and measured using chlorophyll fluorescence, employing the default *Fv/Fm* protocol from a closed FluorCam (Photon Systems Instruments, Czech Republic). The quantification analysis was conducted using the FluorCam 7.0 software.

### Virus-induced gene silencing in *N. benthamiana* and host-induced gene silencing in *P. capsici*

Virus-induced gene-silencing (VIGS) procedure was conducted described to the protocol described by Liu et al. (2002). *A. tumefaciens* suspensions containing pTRV1 and pTRV2 with *SGT1, EDS1, ADR1/NRG1,* and *NRC2/3/4* were mixed at a 1:1 ratio in infiltration buffer (10 mM MES, 10 mM MgCl_2_, and 200 μM Acetosyringone, pH 5.6) to a final optical density (O.D.) at 600 nm of 1.5. This mixture was infiltrated into two leaves of 2-week-old *N. benthamiana* plants. All plants were grown within a walk-in chamber at 24℃ under a 16h/8h (day/night) cycle. After 3 weeks, the upper leaves were utilized for further experiments assessing the efficiency of silencing. Host-induced gene silencing (HIGS) was conducted following the VIGS procedure. After 3 weeks, *P. capsici* strain 40476 was inoculated onto the detached upper leaves. After 6 hours, four leaf discs containing zoospores were collected to evaluate the silencing efficiency.

### Extraction of DNA and RNA and gene expression analysis by qPCR

To assess the biomass of *P. capsici,* total genomic DNA was extracted from four leaf discs adjacent to the *P. capsici*-inoculated area using cetyltrimethylammonium bromide (CTAB) method. For gene expression analysis, total RNA was extracted from four leaf discs using TRIzol reagent (TR118, MRC, USA), and cDNA synthesis was conducted using Superscript II (18064014, Invitrogen, USA) following the manufacturer’s instructions. Quantitative PCR (qPCR) and quantitative reverse-transcription PCR (qRT-PCR) was performed utilizing ExcelTaq™ 2X Q-PCR Master Mix (SYBR, ROX; TQ1110, SMOBIO, Taiwan) with a CFX96 Touch Real-Time PCR Detection System (Bio-Rad, USA). The expression of the PcActin gene was normalized to elongation factor-1a of *N. benthamiana* (NbEF-1a), and the transcript levels were normalized using the internal standard (NbEF-1a). The primer sequences used in this study are provided in Supplemental Table 4.

### Yeast two hybrid assay

Pc12 and truncated Pc12 were cloned into pGBKT7 vector (Takara Bio, Japan), while Rab13-2 and Rab13-2 mutants were inserted into pGADT7 vector. The recombinant plasmids, pGBKT7 and pGADT7, were introduced into yeast strain Y2HGold and Y187, respectively. For yeast two hybrid assay, yeast mating was performed in yeast peptone dextrose adenine (YPDA) medium at 30℃ overnight. The resulting colonies were recovered on a Synthetic Defined medium lacking tryptophan and leucine (SD/-Lue-Trp), and the interactions were validated on SD medium lacking histidine, leucine, and tryptophan (SD/-Lue-Trp-His). The medium plates were incubated at 28℃ and typically photographed in 5 days. For positive and negative controls, commercial yeast constructs were used: positive control (pGBKT7-p53/pGADT7-T) and negative control (pGBKT7-p53/pGADT7-Lam), both provided by Matchmaker™ Gold Yeast Two-Hybrid System (630489, Clontech, USA).

### Co-immunoprecipitation, immunoblot assays, and IP-MS analysis

*N. benthamiana* leaves infiltrated with *Agrobacterium* were sampled at 24–30 hpi for co-IP or western blotting. Total protein was extracted using an extraction buffer (10% [v/v] glycerol, 25 mM Tris–HCl [pH 7.5], 1 mM EDTA, 150 mM NaCl, 1% [w/v] polyvinylpolypyrrolidone, and 1× protease inhibitor cocktail). The extracted proteins were immunoprecipitated with 10 ul of anti-HA magnetic beads (M180-10, MBL, Japan), anti-GFP agarose beads (D153-8, MBL, Japan), or anti-FLAG agarose beads (651502, BioLegend, USA) and incubated for 4 hours or overnight at 4°C. The beads were washed ten times with immunoprecipitation wash buffer (GTEN extraction buffer with 0.015% [v/v] Triton X-100) and resuspended in 10 μl SDS loading dye. Proteins were eluted from the beads by heating at 95°C for 5 min. For western blotting, the immunoprecipitated and input proteins were separated on SDS–PAGE gels and transferred onto PVDF membranes via a Trans-Blot Turbo Transfer System (Bio-Rad, USA). After blocking the membranes with a solution of 5% skim milk prepared in PBST with 0.1% Tween 20, they were incubated with HRP anti-RFP (1:12000, M204-7, MBL, Japan), HRP anti-GFP (1:12000, AB6663, Abcam, UK), HRP anti-HA (1:12000, AB173826, Abcam, UK), or HRP anti-FLAG (1:12000, A8592, Sigma, USA) antibodies at room temperature for 1 h. The membrane was washed twice with PBST for 10 min each before ECL (1705061, Bio-Rad, USA) detection was performed according to the manufacturer’s instructions. For IP-Mass analysis, immunoprecipitated proteins were collected on SDS–PAGE gels and the samples were analyzed using High Resolution LC/MSMS spectrometer (Q Exactive, Thermo Scientific, USA) in NICEM (National Instrumentation Center for Environmental Management, College of Agriculture and Life Sciences, Seoul National University Seoul 151-742, Korea)

### GTPase activity assay

For the GTPase activity assay of Rab13-2, Rab 13-2Q74L, and Rab13-2S29N, E. coli codon-optimized genes were synthesized. Each gene was constructed in a pGEX6-T vector and transformed in BL21 *E. coli* cells to express GST fusion proteins. GPTase activities were measured by GTPase-Glo^TM^ Assay (Promega, V7681). GST-tagged protein samples (1 μg) purified with Pierce® Glutathione Agarose (Thermo, 16100) were used for analysis, and negative controls were constructed by purifying GST. Both samples were incubated with GTPase/GAP buffer containing 10 μM GTP and 1 mM DTT followed by the addition of GTPase-Glo™ reagent. Detection Reagent detects the luminescence of ATP produced from the remaining GTP after hydrolyzing GTP in the reaction. Luminescence was measured using a luminometer (Tecan Spark 10M Multimode Plate Reader), Calculate the GTPase activity as relative luminescence units (RLU). Reaction time was 120 minutes.

### Confocal microscopy

For a confocal microscopy, 5 mm^2^ *N. benthamiana* leaf discs of the infiltrated region were observed 36 hours post *Agrobacteria* infiltration. Images were acquired with an Olympus IX83spinning disk confocal microscope (yokogawa CSW-W1 SoRA) equipped with an ORCA-Fusion Digital CMOS camera under 60 X oil immersion objective (N.A., 1.3) by sequential detection of average of 76 Z stacks. Images were acquired by CellSens Dimension 32 software (Olympus). The 488 nm, and 561 nm laser lines (60% power) were used for EGFP and RFP, respectively. Images were processed using Fiji ImageJ (National Institutes of Health, Bethesda, Maryland, USA) from the maximum Z intensity projections of the confocal images. RFP and moxVenus/GFP were pseudo-colored magenta and green respectively.

### Protein structure prediction

The structures of Pc12 and Rab13-2 were expected using AlphaFold2 in the Google Colab (https://colab.research.google.com/github/sokrypton/ColabFold/blob/main/AlphaFold2.ipynb#scrollTo=svaADwocVdwl). The predicted proteins were visualized using ChimeraX (Pettersen et al. 2021).

### Phylogenetic analysis

The 149 Rab family protein sequences were initially acquired from the *N. benthamiana* genome (v.1.0.1) based on the Ras domain (PF00071) and manually trimmed while considering the Rab description. Alignments were produced using the Multiple Sequence Comparison by Log-Expectation (MUSCLE) software in MEGAX. A phylogenetic tree was constructed using the Maximum Likelihood (ML) method with default parameters and 100 bootstrap replications in MEGAX (Tamura and Nei, 1993; Tamura et al., 2011).

### Statistical analysis

Statistical analyses were performed as described in the figure legends. P values were calculated by Student’s t-test using GraphPad Prism software.

## Data availability

Sequence information of protein-coding genes used in this study was obtained from the FungiDB (https://fungidb.org/), NCBI (https://www.ncbi.nlm.nih.gov/), and Sol Genomics Network (https://solgenomics.net/). The accession numbers for the sequences are as follows: Niben101Scf09596g00001.1 (Rab13-2), Niben101Scf00684g00002.1 (Rab13-3), Niben101Scf03277g02014.1 (Rab13-4), Niben101Scf05709g00001.1 (Rab13-10), Niben101Scf05032g00003.1 (Rab1), Niben101Scf16705g00001.1 (Rab8B), Niben261Chr06g1220001.1 (REP), and Niben261Chr08g0045015.1 (PRA1). All effector sequence information is included in Supplemental Table 1. Rab13-2 (ID:213473) and constitutively active (ID:213474) or inactive mutant (ID:213475), an apoplastic marker (ID: 220490), and a tonoplast marker (ID: 220491) constructs are deposited in Addgene.

## Acknowledgements

This work was supported by grants from National Research Foundation of Korea (NRF) grants funded by the Korean government (MSIT) (NRF-2018R1A5A1023599 [SRC], NRF-2021R1A2B5B03001613, and NRF-2019R1C1C1008698) to D.C. and US National Science Foundation awards (NSF EPS-1655726 for E.P.’s start-up and NSF IOS-2126256 to E.P. and J.C.) and the intramural research program of the U.S. Department of Agriculture, National Institute of Food and Agriculture Hatch Capacity (# 7000762) to E.P. We thank the UWYO INBRE program supported by IDeA from NIGMS, NIH, 2P20GM103432 for providing the spinning disc confocal microscope.

## AUTHOR CONTRIBUTIONS

E.P. and D.C. conceptualized the project. J.Kim, J.Kaleku, J.W., H.J., and. S.J. performed experiments. H.K., M.K., H.J.K., and C.S. provided initial data and materials. J.Kim, J.Kaleku, J.W. and E.P. analyzed the results. J.Kim, E.P., J.W., and D.C. wrote the manuscript. E.P., J.W., and D.C. provided funding to conduct the project.

## Supplemental data

**Supplemental Table 1.**

Pc12 family in *P. capsici* isolates genome

**Supplemental Table 2.**

Candidates targeted by Pc12 in Mass spectrometry analysis

**Supplemental Table 3.**

Rab13-2 interactors from STRING

**Supplemental Table 4.**

Primer sequences used in this study

**Supplemental Figure 1. The expression pattern of Pc12 during the infection phases of *P. capsici.*** *PcHmp1* and *PcNpp1* are utilized as markers for the biotrophic and necrotrophic phases, respectively. Leaf disks were sampled at regular time intervals. Each gene is normalized to *PcTubulin*.

Supplemental Figure 2. Pc12 homologs from *P. capsici* induce cell death, except for PHYCA22034.

(A) Cell death analysis of Pc12 homologs from *P. capsici, P. ramorum, P. cinnamoni,* and *P. sojae.* Pc12 homologs were expressed in 4-week-old *N. benthamiana* and images were photographed at 2 days after agroinfiltration.

(B) The cell death in images (A) was quantified by quantum yield (*Fv/Fm*) using a closed FluorCam system. Data are mean ± SD (n = 9-12). a indicates statistically significant differences by Student t-test (a, P < 0.0001).

**Supplemental Figure 3. Phylogeny tree of Rab family in *N. benthamiana*.**

The red triangle represents the Pc12 target detected in MS, while black triangles represent Rab proteins used in the Co-IP shown in Figure 3B.

**Supplemental Figure 4. Screening of Rab13-2 interactor candidates from STRING database**

Leaves co-expressing Rab13-2 with interactor candidates in Supplemental Table 3 were sampled at 2 dpi, and total protein extracts were subjected to co-IP using anti-GFP agarose beads. (3 experimental repeats)

**Supplemental Figure 5. REP1 and PRA1 prefer to interact the inactive form of Rab13-2 (Rab13-2^S29N^).**

Leaves expressing REP1 or PRA1 with Rab13-2 mutants were sampled at 2 dpi, and total protein extracts were subjected to co-IP using anti-FLAG agarose beads. (3 experimental repeats)

**Supplemental Figure 6. Repetitions of Co-IP assays with REP, Pc12, and Rab13-2 mutants in Figure 4(E)**.

Leaves expressing REP, Pc12 with Rab13-2 mutants were sampled at 12 hpt after 5% ethanol at agrobacteria infiltration, and total protein extracts were subjected to co-IP using anti-GFP agarose beads.

**Supplemental Figure 7. Colocalization of Golgi and ER with Rab13-2 and its mutants in *N. benthamiana*.**

*Cis*-Golgi marker, GO-mCherry, were infiltrated in the epidermal cells of double transgenic plants of RFP-HDEL and GFP-Rab13-2 (A), GFP-Rab13-2^Q74L^ (B), or GFP-Rab13-2^S29N^ (C) *N. benthamiana* plants.

**Supplemental Figure 8. Different impact of Pc12 on secretory pathway and vacuolar targeting pathway.**

(A) Apoplastic marker displacement upon Pc12 expression occurs on 18 hours after the induction (hpi) of Pc12. Interestingly RFP-Pc12 is undetectable under microscope at 18 hpi.

(B) 36 hpi of Pc12 expression, most of observed cells are dead. However, intact vacuolar membranes labeled by tonoplast markers were seldom detected.

